# Large-scale control over collective cell migration using light-controlled epidermal growth factor receptors

**DOI:** 10.1101/2024.05.30.596676

**Authors:** Kevin Suh, Richard Thornton, Payam E. Farahani, Daniel Cohen, Jared Toettcher

## Abstract

Receptor tyrosine kinases (RTKs) are thought to play key roles in coordinating cell movement at single-cell and tissue scales. The recent development of optogenetic tools for controlling RTKs and their downstream signaling pathways suggested these responses may be amenable to engineering-based control for sculpting tissue shape and function. Here, we report that a light-controlled EGF receptor (OptoEGFR) can be deployed in epithelial cell lines for precise, programmable control of long-range tissue movements. We show that in OptoEGFR-expressing tissues, light can drive millimeter-scale cell rearrangements to densify interior regions or produce rapid outgrowth at tissue edges. Light-controlled tissue movements are driven primarily by PI 3-kinase signaling, rather than diffusible signals, tissue contractility, or ERK kinase signaling as seen in other RTK-driven migration contexts. Our study suggests that synthetic, light-controlled RTKs could serve as a powerful platform for controlling cell positions and densities for diverse applications including wound healing and tissue morphogenesis.

## Introduction

Collective cell migration is a fundamental process governing multicellular phenomena such as morphogenesis, wound healing, and cancer invasion^1–3^. The ability to control collective migration – sculpting tissues with high precision using patterned stimuli – could improve our understanding of this fundamental tissue-scale process and serve as a useful substrate for applications ranging from accelerated wound healing to patterning biologically relevant tissue organization.

Over the past decade, various tools have been developed to achieve programmable control over collective cell migration. Tailored ligand gradients can drive chemotactic responses, but programmable control over gradient shape is challenging and requires complex microfabricated devices^4–6^. Micropatterning chemotactic ligands and extracellular matrices can control cellular behavior at high spatial resolution, but these cues typically cannot be dynamically altered once patterned^7–9^. Directing collective migration using electric fields is a promising approach, as electrical cues can drive electrotaxis in many cell types and can be rapidly adjusted in multiple spatial dimensions^10–14^. However, the mechanisms by which cells sense and respond to electric fields are still poorly understood, and precisely manipulating electric fields requires sophisticated device design^10,15^.

Optogenetics also represents a promising approach for guiding cell and tissue motility. Light can be focused precisely in space, rapidly applied/removed, and patterned using simple optical approaches. Moreover, a wealth of light-controlled signaling proteins have been previously developed that could potentially interface with cell motility programs, including light-controlled GTPases and their regulators^16,17^, phosphoinositide 3-kinase (PI3K)^18^, and receptor tyrosine kinases^19^. Exciting work has already demonstrated light-based guidance of individual or small groups of cells^16,17,20–22^ and even morphogenesis in the early *Drosophila* embryo^23,24^, yet optogenetic control of mammalian tissues at macroscopic (millimeter or larger) length scales has not yet been achieved, despite its critical importance for applications ranging from tissue regeneration to organoid production in defined geometries.

We hypothesized that light-gated receptor tyrosine kinases (RTKs) could serve as an ideal platform for achieving optogenetic control over collective cell migration. Receptor tyrosine kinases play essential roles in cell and tissue movement in many different contexts ranging from wound healing and regeneration^25,26^ to developmental collective migration of border cells^27^ and neural crest cells^28^. RTKs also interface with many different potential modulators of cell motility, including Src family kinases^29^, PI 3-kinase^18,30^, and Erk/MAP kinase signaling^20^, enabling them to potentially orchestrate complex downstream programs. RTKs are typically activated by the association of individual receptor molecules upon ligand binding, and multiple groups have developed optogenetic RTK variants based on fusion with protein domains that undergo dimerization or oligomerization upon illumination^19,31–34^. We previously developed two light controlled RTKs – OptoFGFR1 and OptoEGFR – in which the intracellular domains of these receptors are fused to the OptoDroplet protein phase separation system^35^, resulting in rapid, potent, reversible, and spatially controllable activation of either receptor^8,9^.

Here, we report that our OptoEGFR system can be used to drive large-scale, light-controlled collective migration of mammalian cells. We observe distinct effects of OptoEGFR stimulation on collective migration depending on the geometry of the tissue and illumination pattern. Tissue densification was produced when a local light input applied to an interior region within a continuous monolayer, driven by converging cell movement into the illuminated region. Conversely, illumination of a tissue edge drove rapid tissue expansion at speeds ∼40% faster than un-illuminated control tissues. We also observed an overall increase in tissue motility and outward migration speed when tissues were globally illuminated. Overall, these data suggest that OptoEGFR can both act as a local directional cue to guide collective migration, and as an overall amplifier of directional cell movement initiated by other non-optogenetic sources.

Pharmacological perturbations and tissue patterning experiments revealed that large-scale tissue movements were primarily driven by physical interactions between cells, not diffusible ligand gradients; that ERK signaling and myosin-driven contractility were dispensable for tissue movement; and that PI3K signaling activity was required for the effect. Our data is consistent with a model where boundaries of the light pattern drive directional tissue flows, a principle that can be used to guide tissue patterning into more complex structures.

## Results

### OptoEGFR stimulation triggers both local tissue convergence and enhanced outgrowth

We initially characterized the cell motility effects of OptoEGFR and OptoFGFR light-controlled receptor tyrosine kinases^32,33^ (**Fig. 1A-B**). In each case, the intracellular domains of the receptor tyrosine kinases was fused to the FusionRed fluorescent protein as well as the membrane OptoDroplet system^35^, which is composed of a myristoylation tag to drive membrane localization, the FUS disordered N-terminal sequence, and the Cry2^PHR^ domain which undergoes oligomerization upon illumination with 450 nm light^36,37^.

**Figure 1.**
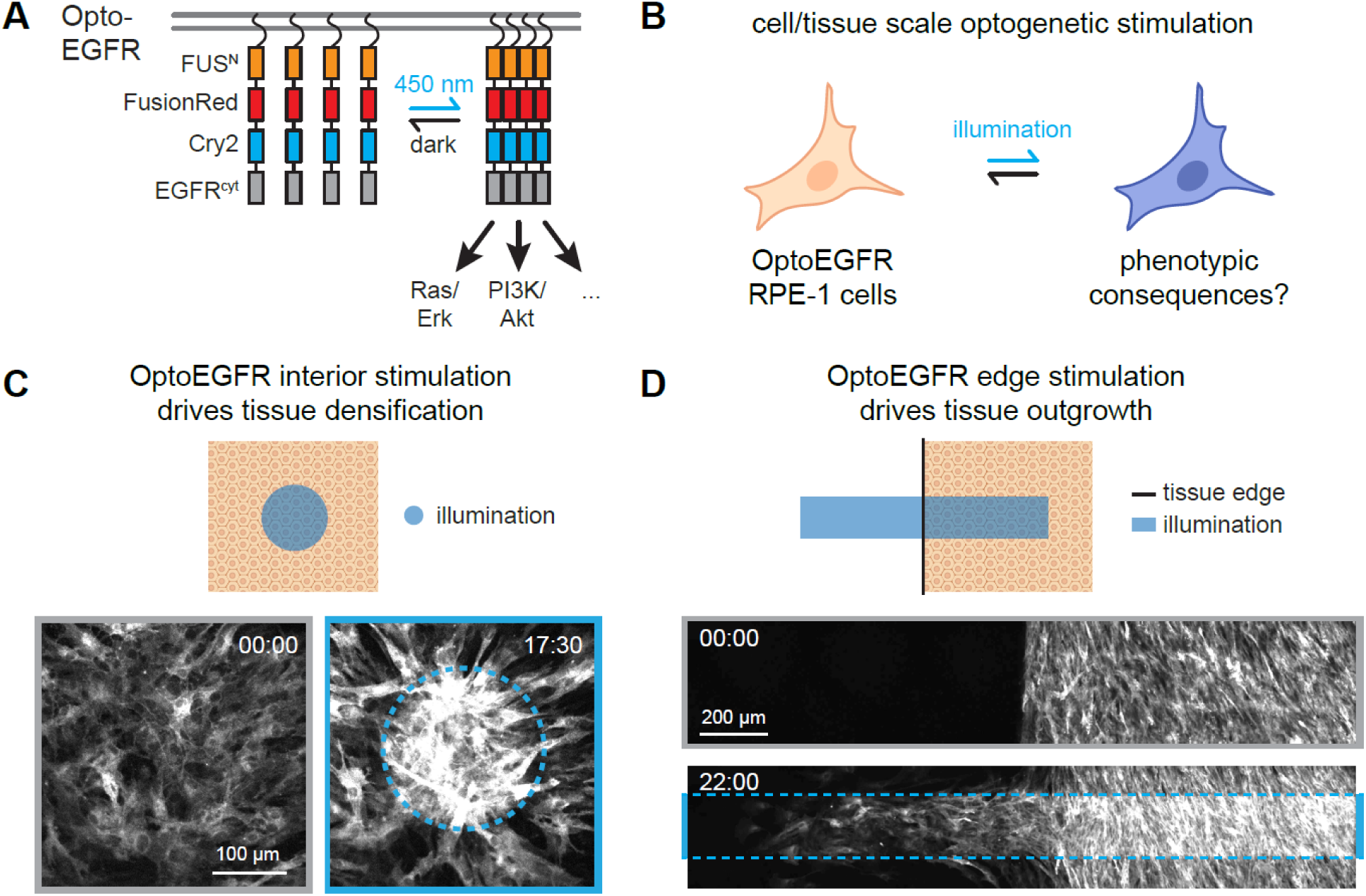
Optogenetic EGFR stimulation directs tissue movement. (**A**) Schematic of OptoEGFR construct for blue light-inducible EGFR clustering and signaling pathway activation. The system fuses an N-terminal membrane localization tag to the OptoDroplet system for light-inducible clustering (FUS^N^-FusionRed-Cry2) and the cytosolic domains of EGFR. (**B**) We set out to assay for phenotypic consequences to OptoEGFR stimulation. (**C-D**) Light stimulation of OptoEGFR RPE-1 cells produces dramatic tissue movements, including tissue densification in illuminated interior regions (in **C**) and tissue outgrowth from illuminated tissue edges (in **D**). Images show FusionRed channel; stimulation time shown

We introduced OptoEGFR or OptoFGFR using lentiviral transduction into RPE-1 cells, an immortalized human retinal pigmental epithelial cell line^38^, plated confluent monolayers of each cell line, and locally stimulated them with pulses of 450 nm blue light delivered every 2-3 min using a digital micromirror device on our microscope at an intensity of 65 mW/cm^2^. We imaged cells in the FusionRed channel, which marks OptoEGFR expression and localization. We found that illumination elicited profound changes in tissue organization, with OptoEGFR cells undergoing rapid and sustained movement into the illuminated region (**Fig. 1C**; **Video S1**). We further observed that local illumination at the edge of an OptoEGFR-expressing tissue produced a distinct effect, with cells rapidly moving outward from the edge to fill the illuminated region (**Fig. 1D**; **Video S2**).

Light-induced migration phenotypes were not a general feature of optogenetic receptor tyrosine kinase activation. Rather than rapid light-induced migration into illuminated regions, we found that OptoFGFR-expressing cells were gradually excluded from the illumination region (**Fig. S1A**), consistent with our prior observations of retraction away from illuminated regions in individual OptoFGFR-expressing NIH3T3 mouse fibroblasts^32^. We observed similar degrees of ERK phosphorylation with both OptoEGFR and OptoFGFR-expressing cells (**Fig. S1B**), suggesting that this difference in cellular responses was not driven by an absolute difference in receptor activity but rather different intracellular signaling pathways engaged by the two receptors.

We also observed similar OptoEGFR-driven tissue movement in a second human cell line, MCF10A breast epithelial cells expressing the ErkKTR biosensor for Erk mitogen-activated protein kinase (MAPK) activity^39^. Illumination drove rapid export of the ErkKTR from cells only within the illuminated region as well as tissue convergence in an analogous manner to what was observed in RPE-1 cells (**Fig. S1C**; **Video S3**). These data confirm that illumination drives localized OptoEGFR activation, and that the light-induced tissue movement triggered by OptoEGFR generalizes across multiple cellular contexts.

Our migration data present an apparent paradox because the same optogenetic tool and light stimulus can drive opposing effects: either convergent motion and an increase in cell density when illumination is applied at interior positions, or divergent outgrowth and expansion from a tissue edge. In subsequent experiments, we sought to quantify both types of motion and to dissect the basis for light-induced tissue movement to resolve this paradox.

### OptoEGFR tissue densification is driven by collective migration at the illumination boundary

We next sought to better understand and quantify how OptoEGFR stimulation drives convergent motion in a confluent monolayer. Local stimuli can often elicit global responses in collective systems^40,41^, so we sought to scale up our stimulus and imaging conditions to the mm-cm scale in living tissues. Collective cell behaviors depend heavily on tissue size and shape^42^, so we first engineered precise arrays of replicates of 6-mm diameter circular tissues using our tissue stenciling approach^43^ to increase throughput, improve statistical power, and ensure directly comparable tissues. Typically, localized optogenetic stimuli are applied to cells and tissues using digital micromirror devices through the imaging light path, which restricts patterned stimuli to a single field of view. To expand optical stimulation to a larger length scale, we instead projected various illumination patterns through the transmitted light path using laser cut photomasks (see **Methods**) placed directly in the light path of the condenser of an inverted microscope, which allowed us to illuminate OptoEGFR-expressing RPE-1 cells over centimeter length scales (**Fig. S2A-B**). We then programmed an automated microscope to uniquely align each of the large tissues with specific patterns on the photomask to allow us to use one photomask to stimulate multiple tissues (see **Methods**).

We used this approach to apply circular illumination patterns with 200 μm, 1 mm, and 2 mm diameters at the center of confined, 6 mm diameter RPE-1 tissues and imaged cells in the FusionRed channel (**Fig. 2A**; **Video S4**). Light stimulation drove rapid tissue movement into the boundary of the illuminated region that gradually filled in toward the center. Outside the illumination boundary, a broad region of decreased cell density was also observed, suggesting that cells were displaced from hundreds of micrometers away from the surrounding tissue into the illuminated region, indicating a large correlation length. We quantified the converging migratory behavior of the illuminated tissue with particle image velocimetry (PIV) analysis on the time-lapse images (**Fig. 2B**) to produce spatial maps of migration dynamics. Confirming our qualitative observations, a local velocity vector map showed strong converging motion generated at the illumination boundary and nearby un-illuminated tissue, oriented toward the illumination center (**Fig. S2C**). A kymograph of the radial component of tissue velocity revealed that the convergent motion was relatively stable over time, extending ∼500 μm from the illumination boundary (**Fig. 2B**).

**Figure 2.**
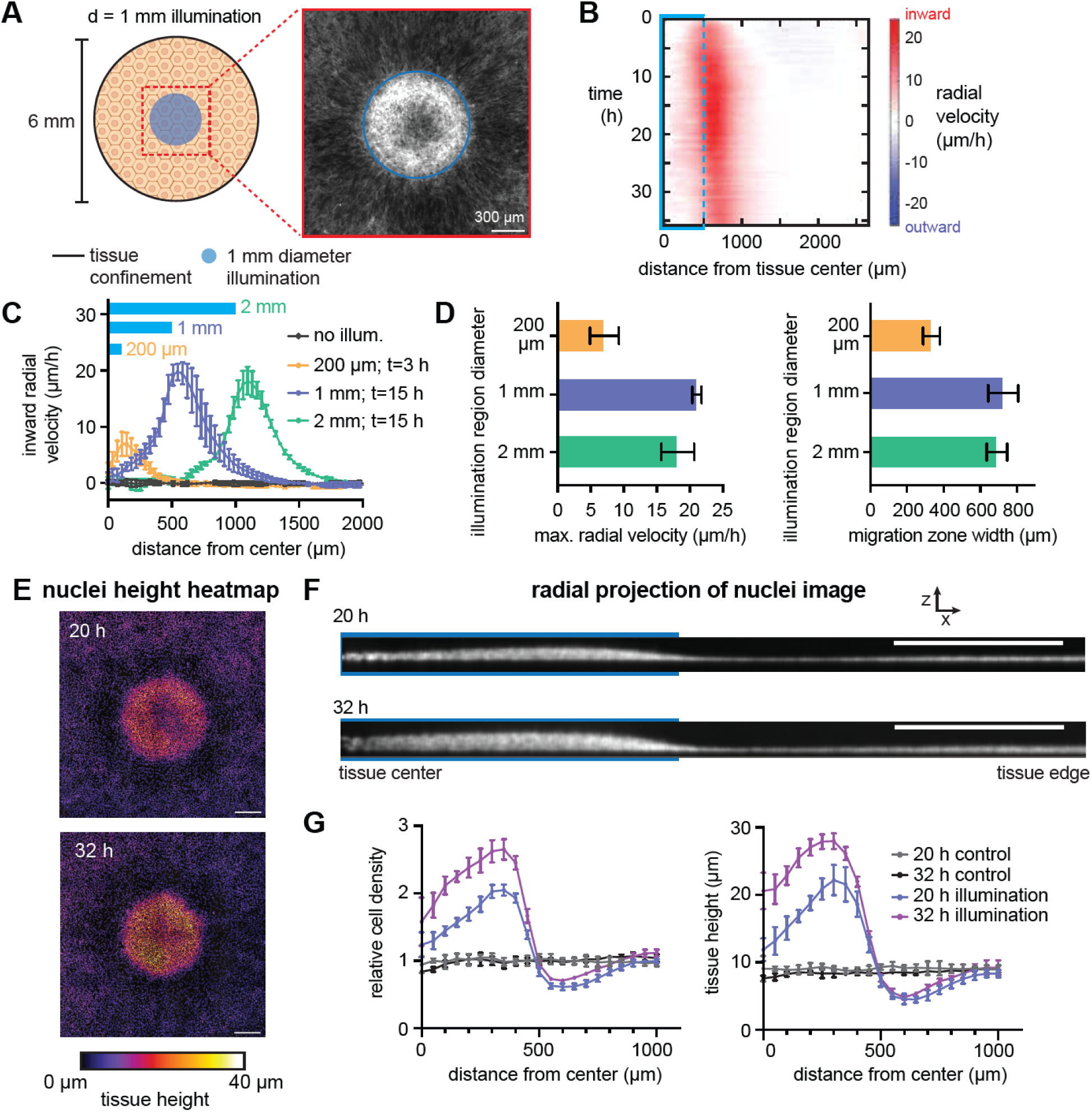
Quantifying large-scale tissue densification induced by OptoEGFR stimulation. (**A**) Schematic of large-scale tissue stimulation assay. Circular light stimuli of varying sizes are applied to the center of a confined 6-mm RPE-1 tissue and cell movements are imaged using the membrane FusionRed tag. (**B**) Quantification of tissue velocities as a function of time and distance from the tissue center. The illumination boundary is shown (blue region). (**C**) Quantification of tissue velocity as a function of position at the fastest-moving time point for 200 μm, 1 mm, and 2 mm-diameter illumination patterns. (**D**) Maximum radial velocity and migration zone width corresponding to the curves in **C**. N=9,3,3,3 for control and each illumination pattern, respectively. (**E**) Confocal stacks of nuclei staining for 1 mm illumination pattern, colored by tissue height at the indicated times after illumination. Scale bar: 300 μm. (**F**) Sum-projection along the radial coordinate for tissues in **E**. (**G**) Quantification of data in **F** showing relative cell density and tissue height as a function of radial position. N=3,5 for control and illuminated

We next quantified the spatial profile of tissue velocities for each illumination pattern (**Fig. 2C**), focusing on the time point at which the maximum velocity was achieved for each illumination pattern (3 h for 200 μm tissue; 15 h for 1- and 2-mm tissues) (**Fig. S2D-E**). We observed a sharp, relatively symmetric peak in tissue velocity near the border of the illumination area, with slower movement farther into the illuminated region or in the un-illuminated exterior region (**Fig. 2C**). Tissue movement was fastest and most sustained for millimeter-scale illumination patterns, which produced a ∼3-fold higher peak velocity compared to the 200 μm illumination pattern (**Fig. 2D**). These data suggest that localized optogenetic stimulation could be well suited for driving tissue reorganization even at macroscopic length scales.

Notably, the zone in which directional migration was observed was confined to a region near the illumination boundary of similar width for both the 1 mm and 2 mm illumination pattern (**Fig. 2D**). For the 2 mm diameter pattern, cells within the illuminated region more than 1 mm from the light boundary did not undergo substantial movement, despite consistent illumination. These data suggest that light-induced collective migration is confined near the interface between illuminated and un-illuminated tissues.

We also systematically varied illumination dose for a fixed geometry to test how the strength of OptoEGFR activation alters migratory responses (**Fig. S2F**). Compared to our base case (5 sec per min of 450 nm light exposure), we found that increasing the illumination frequency by three-fold (5 sec every 20 sec) dramatically decreased overall tissue migration, whereas a lower illumination dose (4 sec every 2.5 min) modestly decreased movement speed. These data suggest that the extent of tissue migration varies with both the illumination pattern geometry and the illumination schedule of OptoEGFR activation, and might reflect an optimal, intermediate level of RTK activity for driving cell migration or competing timescales for light-induced changes in cell/tissue mechanics.

If cells migrate over long distances to enter regions of light-induced OptoEGFR activity, we might expect a dramatic increase in cell density or a transition from 2-dimensional to 3-dimensional tissue organization over time. Indeed, we found that light-induced collective migration also led to pronounced tissue thickening that was evident in confocal z-stacks of the illuminated tissue (**Fig. 2E**) as well as radial profiles obtained by summing across radial slices of the nuclear intensity image (**Fig. 2F**). Both tissue height and cell density rose sharply to a peak ∼50 μm interior to the illumination boundary to values approximately 3-fold higher than un-illuminated control tissues (**Fig. 2G**). We also observed depletion of cells outside the illumination area, consistent with the elongated morphology of cells just outside the illumination area (**Fig. 1C**). We also found that light-induced tissue densification persisted for at least 40 h after a shift back to darkness (**Fig. S2G-H**), consistent with a model where localized light stimuli drive irreversible cell rearrangements and permanent changes to tissue structure. Taken together, these data demonstrate that optogenetic EGFR stimulation drives rapid collective migration toward sites of illumination, leading to millimeter-scale changes in tissue organization.

### Global tissue illumination drives tissue fluidization and rapid outgrowth

We next set out to measure the effect of OptoEGFR stimulation at tissue boundaries. We reasoned that local illumination at a tissue edge would combine two effects: light-induced edge outgrowth and light-induced convergence at interior boundaries. To simplify the geometry, we reasoned that globally applying light to unconfined tissues would not produce a light boundary within the tissue and thus avoid regions of local tissue convergence. We grew 2-mm diameter circular tissues in a confining stencil, allowing them to reach confluency, and then removed the stencil and monitored tissue outgrowth for 24 h in the presence or absence of global 450 nm illumination (**Fig. 3A**, **Video S5**). We found that illuminated tissues indeed exhibited more rapid outgrowth compared to unilluminated controls, with movement that extended deeper within the interior of the tissue (**Video S5**). We quantified the edge expansion by fitting the expanding tissue to a circle at each time point (**Fig. 3B**) and estimated the speed of outgrowth from the rate of radial growth (**Fig. 3C**). We found that illuminated tissues grew consistently faster than their unilluminated counterparts, with a ∼40% increased speed on average over the time course (**Fig. 3D**). These data indicate that OptoEGFR stimulation can exert global effects on tissue movement, increasing the rate of expansion of unconfined tissue.

**Figure 3.**
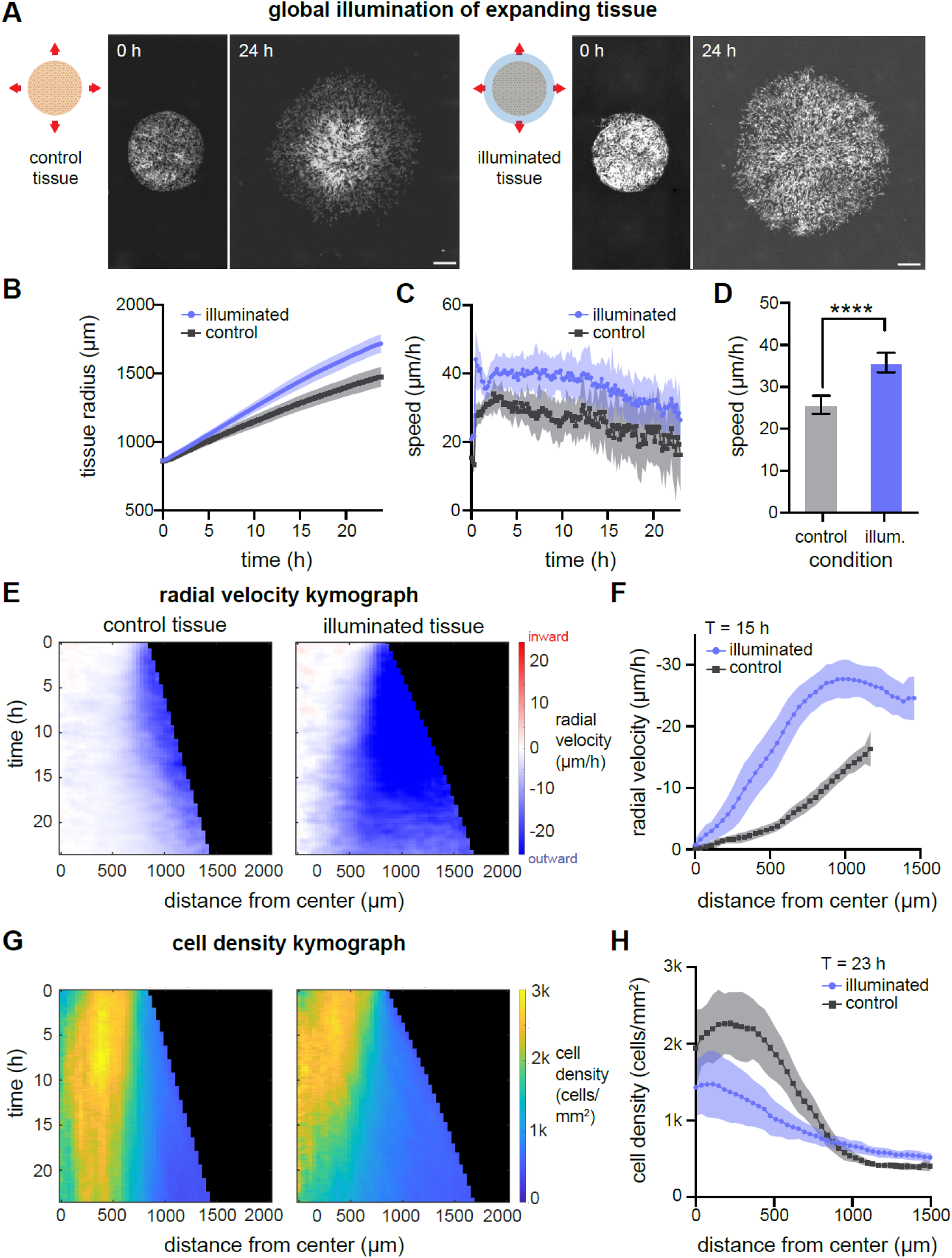
Global illumination drives tissue fluidization and enhanced outgrowth. (**A**) Initial and final (24 h) images of OptoEGFR RPE-1 tissues in darkness or under global 450 nm illumination. Scale bar: 500 μm. (**B-C**) Quantification of tissue radius over time (in **B**) and radial tissue velocity computed as the rate of change of tissue radius growth (in **C**). (**D**) Mean edge speed compared between control and globally illuminated tissue. N=15,18 for illuminated and control tissues, respectively. (**E**) Kymograph of radial velocity as a function of position and time, measured by particle image velocimetry on expanding tissues. (**F**) Quantification of radial velocity as a function of position at 15 h after barrier removal. N=11,15 for illuminated and control tissues, respectively. (**G**) Kymograph of cell density as a function of position and time for illuminated and unilluminated expanding tissues. (**H**) Quantification of cell density as a function of position at 23 h after barrier removal. N=11,15 for illuminated and control tissues, respectively.

We notice that globally illuminated OptoEGFR tissues exhibited increased collective motion not only at tissue edges but also at interior regions (**Video S5**). To quantify this effect, we mapped the local velocity field of the entire tissue using PIV analysis (**Fig. 3E**). In the control case, outward tissue flow was confined to the outermost ∼500 μm, with minimal movement at interior positions (**Fig. 3E**, blue region). This observation is consistent with prior studies of expanding tissue monolayers^42,44,45^ as well as the prior observation of decreased movement at high cell densities termed contact inhibition of locomotion (CIL)^46–48^. Tissue-scale CIL has been interpreted as a jamming transition^49,50^ that coincides with high cell density^46,51,52^. In contrast, interior regions of illuminated OptoEGFR-RPE-1 tissues gradually began to flow outward (**Fig. 3E**). Strikingly, illuminated tissues maintained outgrowth speeds at interior positions that were even higher than the peak speeds observed at the periphery of control tissues (**Fig. 3F**). This may indicate a general fluidization of the bulk as the previously ‘solid-like’ interior gave way to increased motility.

To better characterize the interplay between migration and cell density, we used the Hoechst Janelia Fluor 646 live-cell dye to stain cell nuclei and monitor cell density throughout the tissue (**Fig. 3G-H**). Despite initially similar density profiles, illuminated tissues gradually decreased in cell density in coordination with increased outward tissue speed, whereas control tissues retained the high-density interior that is usually observed epithelial monolayer expansion^42,43^. In summary, global illumination of dense tissues with free edges enhanced the tissue’s outgrowth and promoted fluidization of interior regions. The increase in cell movement and decrease in cell density of illuminated tissues is reminiscent of epithelial tissue unjamming, the transition of tissue phase from static solid-like phase to motile fluid-like phase^53,54^.

### Light-induced tissue movement depends on cell-cell contact and PI 3-kinase signaling

What processes downstream of OptoEGFR stimulation drive collective migration? Many cellular processes have been implicated in RTK-directed cell migration. Recent studies in Madin Darby canine kidney (MDCK) cells provide evidence of EGFR-related collective cell migration being driven by a feedback loop between intracellular Erk kinase activity, cell contractility, and ADAM17-triggered shedding of epidermal growth factor (EGF) to stimulate Erk activity neighboring cells^20,21^, and prior studies also implicate myosin contractility initiated by Rho kinase (ROCK)^55^ and PI 3-kinase activity^18,22,56^ as RTK-dependent drivers of cell motility. We thus sought to dissect which cellular processes were responsible for the profound light-induced tissue reorganization that we observed.

We first set out to identify the basic principles governing large-scale cell movement in our system. Our prior experiments revealed oriented cell movement towards the light input that appeared to be restricted to 1 mm region centered on the illumination boundary (**Fig. 2C**). How does the illumination boundary drive tissue movement, and how can a localized light stimulus produce effects hundreds of microns away? We considered two broad classes of tissue level coordination: diffusion of ligands from illuminated to un-illuminated regions, and mechanical coupling, where movement of illuminated cells is sensed through cell-cell contacts or changes in cell density^57,58^.

To discriminate between these broad classes of models, we designed an experiment to determine if differences in illumination could be transmitted across a physical discontinuity between two tissues^59^ (**Fig. 4A**; **Video S6**). We seeded two identical tissues separated by a 300 μm gap, a distance which was shorter than the light-induced migration zone produced in the un-illuminated region of a continuous tissue (**Fig. 2D**). We then illuminated the right-hand tissue and monitored the outgrowth speed of the unilluminated left-hand tissue, comparing outgrowth between the tissue edges that were proximal and distal to the illuminated tissue (**Fig. 4A**).

**Figure 4.**
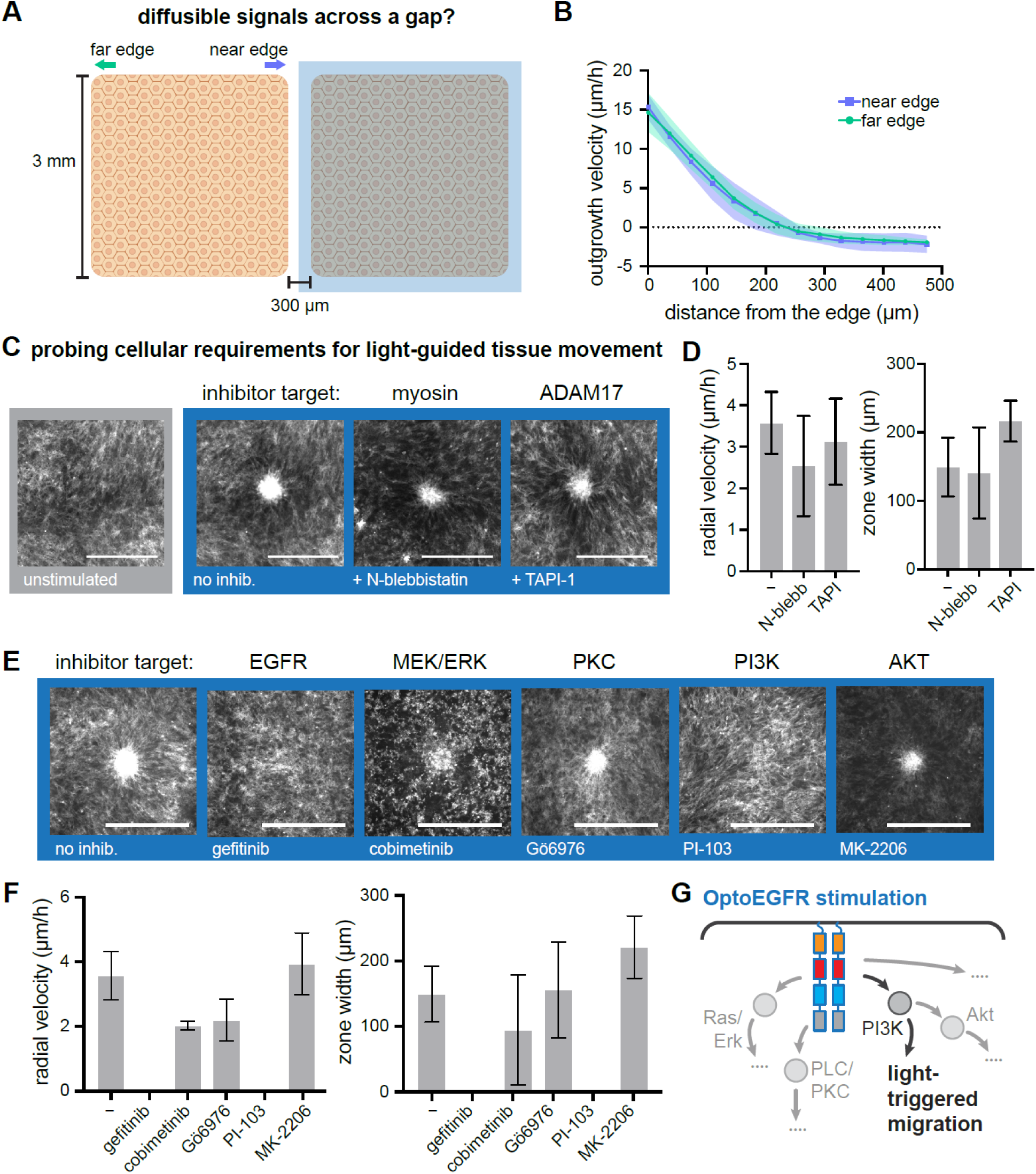
Interrogating the molecular basis for light-induced collective cell migration. (**A**) Schematic of experiment to test for the role of diffusible signaling and an illumination boundary on cell movement. Two tissues are plated with a 300 μm gap and allowed to freely expand while one tissue is illuminated. (**B**) Outgrowth rates as a function of distance from tissue boundary at the near and far edges of the unilluminated tissue in A. N=8 tissues across 2 experiments. (**C**) Images of tissues stimulated as in Fig. 2 with a 200 μm illumination circle in the presence of the indicated chemical inhibitors. Scale bar: 500 μm. (**D**) Quantification of peak radial velocity (left) and migration zone width (right) for illuminated tissues treated with each compound. N=6,6,5 tissues across 2 experiments for control, N-blebbistatin, and TAPI-1 respectively. (**E**) Images of tissues stimulated as in **C** in the presence of the indicated chemicalinhibitors. Scale bar: 500 μm. (**F**) Quantification of peak radial velocity (left) and migration zone width (right) for tissues stimulated as in **E**. N=6,5,3,3,3,5 for conditions labeled left to right. (**G**) Schematic of EGFR signaling and inferred control over light-induced cell migration.

Quantification of tissue outgrowth revealed no difference in outgrowth speeds at the near and far edges (**Fig. 4B**). No large-scale migration was observed toward the illuminated region, suggesting that directed migration requires the projection of a light-dark boundary on cells, not in the gap between cells.

We also performed independent experiments to specifically test for roles of EGF ligand release and cell contractility in tissue-scale motility, as suggested in recent work^20^. We prepared confluent 3 mm-diameter tissues and illuminated a 100 μm-diameter central region to induce local tissue densification in the presence of the ADAM17 inhibitor TAPI-1 or the contractility inhibitor N-blebbistatin (a non-photosensitive variant of the classic myosin inhibitor blebbistatin). Neither N-blebbistatin nor TAPI-1 treatment blocked light-induced tissue densification or long-range cellular movements (**Fig. 4C**; **Video S7**), and tissue movements reached similar peak velocities in all three cases (**Fig. 4D**). However, we note that N-blebbistatin treatment appeared to broaden the migration zone deeper into the unilluminated tissue, consistent with prior observations in MDCK cells that blebbistatin can reduce cell-cell friction and lead to larger regions of coordinated migration^60^ (**Fig. 4D**, right). Overall, these results suggest that diffusible ligand stimulation is dispensable for large-scale tissue movements downstream of OptoEGFR, and that cell-cell contact is required for transmission of information between regions of local OptoEGFR activation and neighboring un-illuminated tissues.

To gain further insight into the signaling pathways involved in coordinating OptoEGFR-induced cell movement, we again performed the light-induced migration assay of **Fig. 4C** in the presence of kinase inhibitors directed at key nodes in the EGFR pathway: EGFR, PI3K, AKT, MEK, and PKC (**Fig. 4E**; **Video S7**). As expected, we found that the EGFR inhibitor gefitinib completely prevented light-induced tissue movement (**Fig. 4E-F**). In contrast, cells retained strong light-induced movement in the presence of the MEK inhibitor cobimetinib, the PKC inhibitor Gö6976, and the Akt inhibitor MK-2206. We observed substantial cell death throughout the tissue during 24 h incubation with the MEK inhibitor cobimetinib, consistent with the importance of mitogen-activated protein kinase (MAPK) signaling for long-term cell survival. Consistent with the dispensability of MEK/Erk signaling for tissue movement in this system, we found that MCF10A cells expressing OptoSOS, an optogenetic system to directly activate Ras/ERK signaling downstream of RTKs^61,62^, had no effect on tissue movement in MCF10A cells also expressing the ErkKTR biosensor, despite similar Erk activation within the illuminated region in both cases (**Video S8**). In contrast, the PI 3-kinase inhibitor PI-103 was the only downstream inhibitor tested to completely block light-induced migration of OptoEGFR RPE-1 cells, phenocopying receptor inhibition by gefitinib. This is also consistent with the role of PI 3-kinase in directed cell migration and emphasizes that optoEGFR activation is likely to act directly at the level of front-rear cell polarity^63^, rather than purely ‘pulling’ cells along by contraction. Similar inhibitor results were also obtained in OptoEGFR MCF10A cells (**Fig. S3**), suggesting that the mechanisms underlying OptoEGFR-induced cell movements are general across cellular contexts.

### Illumination boundaries provide directional information to sculpt tissue organization

Taken together, our results suggest a model for how OptoEGFR stimulation drive tissue movements in both illuminated and un-illuminated regions (**Fig. 5A**). At the illumination boundary, partial illumination of individual cells triggers localized activation of EGFR and its downstream effector PI 3-kinase, leading to cell movement into the illumination region.

**Figure 5.**
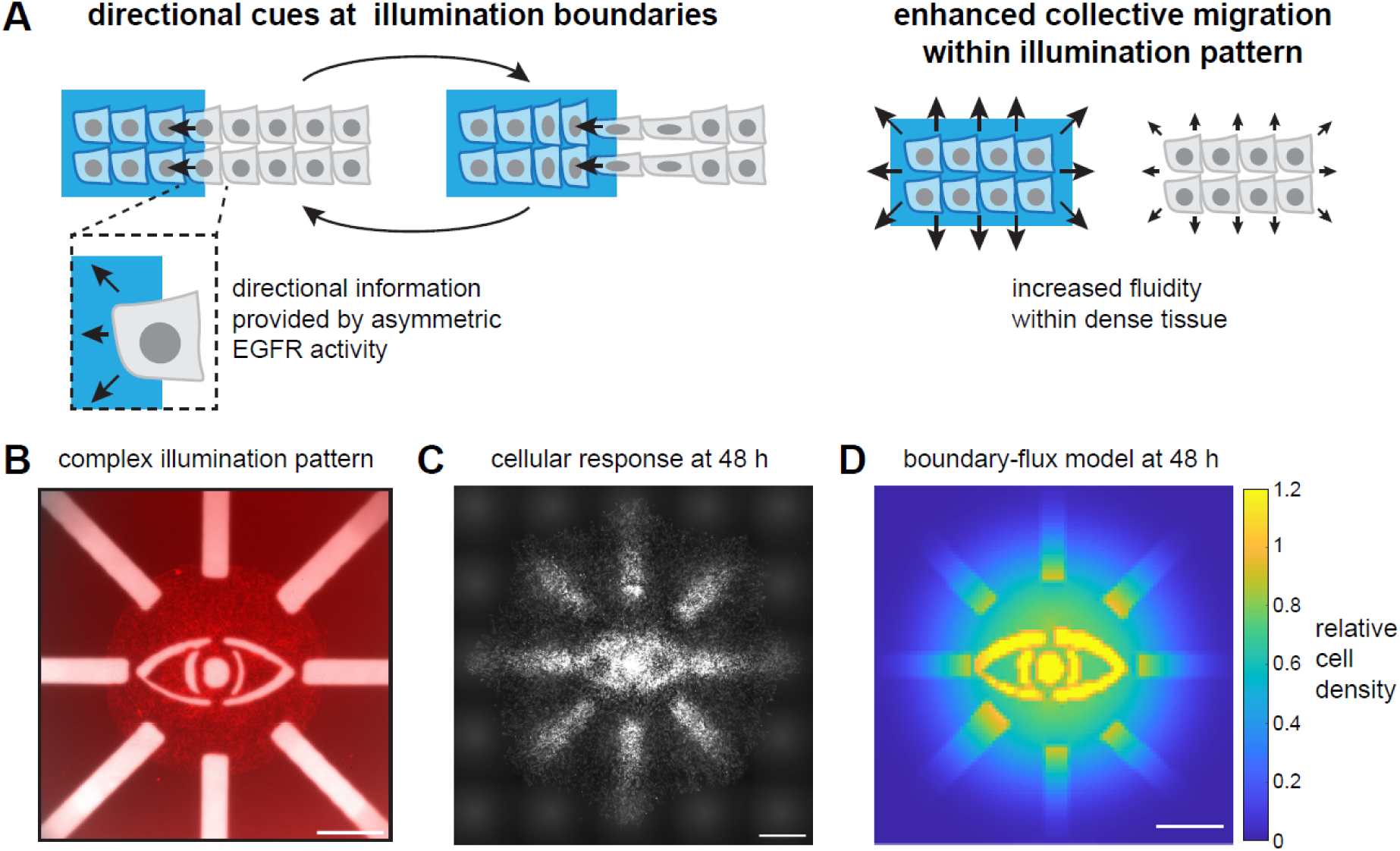
Local and global cues drive OptoEGFR-induced light-induced collective cell migration. **(A)** Conceptual model for how different illuminattion geometries affect collective cell movement. Left: Partially illuminated cells experience a directional cue mediated by PI 3-kinase, driving movement into the illuminated region. This movement can exert force on neighboring un-illuminated cells or leave a lower-density gap to drive these neighboring cells’ movement into the light, repeating the cycle. Right: whole-cell illumination produced by global light stimuli drives distinct effects, including increased tissue fluidity and more rapid outgrowth velocities. (**B**) Driving complex tissue patterning with a combination of interior and edge illumination patterns. Left: illumination pattern applied to a circular tissue; Middle: FusionRed fluorescence imaging after 48 h of illumination and expansion; Right: a simple mathematical model implementing tissue flux at illumination boundaries and diffusion captures qualitative features of the tissue pattern. Scale bar: 1 mm.

Consistent with this picture, optogenetic PI 3-kinase stimulation has been observed to act as a directional cue to guide motility of individual cells^18,22^. Nearby un-illuminated cells would then move toward the illumination boundary, either through forces applied to cell-cell contacts or to fill the gap left by their neighbor at the illumination boundary, leading them to be partially illuminated and repeating the process. Our data also indicates that a second set of phenomena modulate cell movement within illuminated regions, where OptoEGFR stimulation increases both tissue fluidity and edge outgrowth speed (**Fig. 5A**; **Fig. 3A**).

To gain confidence in this conceptual model, we set out to test whether it would be sufficient to recapitulate arbitrary, complex patterns of light-controlled cell movement. We thus implemented a simple mathematical model of light-controlled tissue flows that could be simulated on the same geometry as our experiments. The model assumed a continuous tissue with two sources of tissue movement (1) outward diffusion into un-occupied space (using an effective diffusion parameter *D*, and (2) cell flux at illumination boundaries into the illuminated region (using a boundary flux parameter *k*). The two model parameters *D*=50 μm^2^/min and *k*=1 min^-1^ were qualitatively estimated from the observed rates of tissue outgrowth and light-induced movement throughout our experiments (see **Methods**; **Fig. S4**). Importantly, our continuum model is meant to be a qualitative, simplified implementation to investigate the consequences of a minimal set of biological assumptions (cell flux at illumination boundaries) and does not capture more complex tissue features such as density-driven jamming or light-induced tissue fluidization.

We generated a complex illumination pattern that incorporates multiple domains of tissue densification and outgrowth (**Fig. 5B**) and applied it to both an OptoEGFR RPE-1 monolayer and our mathematical model. We observed the evolution of a complex 3-dimensional tissue structure, with regions of light-induced tissue densification at regions of interior illumination, as well as enhanced outgrowth from illumination at tissue edges (**Fig. 5C**; **Movie S9**). This pattern was qualitatively matched by model simulations, where tissue densification was driven by flux of cells into the illuminated region, and enhanced outgrowth at the tissue boundary resulted from the higher cell density produced by this cell influx. We conclude that tissue flux at illumination boundaries is a simple principle that is sufficient to explain many light-induced tissue movements and is likely to be useful as a starting point for sculpting complex tissue architectures with more sophisticated illumination protocols.

## Discussion

Optogenetics is a powerful tool for spatiotemporal control of cellular behavior. Building upon previous studies focused on subcellular level control^21,33,35^, we investigated the possibility of macroscopic, tissue-level behavioral control using light illumination. Illumination of millimeter-scale OptoEGFR expressing RPE tissue induced two profound phenotypes for tissue-scale movement: (1) tissue densification into local regions of illumination, and (2) accelerated outgrowth at tissue edges. These phenotypes might initially seem contradictory, as the same stimulus (blue light) and cellular context (OptoEGFR cells) can either trigger formation of high-density domains within a tissue or expand outward to low density, depending on the geometry of the tissue and light pattern.

To study tissue flows into local regions of illumination, we projected circular illumination patterns to the inside of the tissue resulted in tissue movement that was distributed over ∼1 mm, with a peak speed at the illumination boundary. The extent of the collective migration was dependent on the size of the illumination pattern, with larger migration speeds and more sustained movement obtained with larger stimulation patterns (**Fig. 2C-D**). This phenomenon might be explained by the high cell densities that are reached in the small illumination patterns, which have a high perimeter but low area, thus driving tissue flow into a region of limited size. We also observed that there is an optimal intermediate light dose for driving tissue migration (**Fig. S2F-G**), with more frequent illumination dramatically inhibiting tissue flows. It is counterintuitive that higher light doses did not trigger a more potent migratory response, which might be explained through dose-dependent effects of EGFR signaling on downstream signaling or motility programs.

To study the effects of OptoEGFR illumination on tissue outgrowth, we monitored expansion of tissues under global illumination and found that the outgrowth of the tissue could be accelerated by ∼40% (**Fig. 3A-D**). Illumination not only affected the edge of the tissue but also enabled free edge expansion to propagate deeper into the tissue, decreasing cell density at the tissue center (**Fig. 3E-H**). Our findings are reminiscent of the solid-like to fluid-like tissue phase transition termed un-jamming that has been reported in other epithelial contexts^53,54^. We note that our RPE-1 cell line’s tissue architecture exhibited spider web-like tissue structure, not a classic epithelial geometry with tightly packed configuration cell bodies filling the entire space, where morphology of each cell (shape index) is used to determine the phase of the tissue based on the energy barrier for cellular junction restructuring^50,64^. Further study would be necessary to determine whether the formal concepts of tissue unjamming can be translated to light-induced fluidization observed in our system.

We propose a simple model to resolve the apparent contradiction of densification at interior regions and outgrowth at cell boundaries (**Fig. 5A**). At the illumination boundary, cells that are partially illuminated migrate directionally toward the illuminated region. Their un-illuminated neighbors then enter the illumination boundary, either by being pulled along cell-cell contacts or by migration into the lower-density region produced by their neighbor’s movement. These cells are now exposed to a partial light stimulus and the process repeats. Consistent with this model, we find that cells must be present at the illumination boundary for directional migration into the illuminated region to occur (**Fig. 4A**). Importantly, cells that are wholly illuminated do not experience a directional cue and are free to expand in any direction, including outward from a tissue edge.

Our study also sheds light on the essential molecular mechanisms for RTK-driven tissue flows. Combining light stimulation with small-molecule inhibitor treatment reveals that PI 3-kinase signaling is essential for light-induced collective cell migration, emphasizing that OptoEGFR acts at the level of cell direction-sensing, producing a collective migration polarity. Our data suggest that OptoEGFR-driven cell migration operates *via* distinct principles from those suggested in recent studies of EGFR-driven cell movement in MDCK epithelial monolayers. Our experiments suggest that neither ADAM17 activity nor EGF ligands diffusion is required to coordinate tissue-scale cell movements downstream of OptoEGFR stimulation, and we further find that Erk activity is neither necessary (using MEK inhibitor treatment) nor sufficient (using OptoSOS stimulation) for light-induced tissue movement in either RPE-1 or MCF10A cells. These data suggest that RTKs can trigger cell movement through a variety of distinct intracellular mechanisms depending on cellular context, and MDCK collective cell migration may represent a distinct mode from the cell lines studied here.

Overall, our study demonstrates that light-controlled tissue movement represents a powerful and controllable means to drive tissue rearrangements, which could find utility in applications where tissue organization is disrupted such as wound healing, tissue regeneration, and restoring proper tissue organization in cases of developmental disorders.

## Methods

### Experimental model and subject details

#### Cell culture

RPE cells were cultured in DMEM/F12 (Gibco,11320033) supplemented with 10% fetal bovine serum (R&D Systems, 26140079), 1% L-glutamine (Gibco,25030081), and 1% penicillin/streptomycin (Gibco,15140122). MCF10A-5E cells^70^ were cultured in DMEM/F12 supplemented with 5% horse serum (Invitrogen,16050122), 20 ng/mL EGF (Peprotech, AF-100-15-1MG), 0.5 µg/mL hydrocortisone (Sigma-Aldrich,H0888), 100 ng/mL cholera toxin (Sigma-Aldrich,C8052), 10 µg/mL insulin (Sigma-Aldrich), and 1% penicillin/streptomycin. All cells were maintained at 37°C and 5% CO2. Cells were tested to confirm the absence of mycoplasma contamination.

### Method details

#### Plasmid construction

All constructs were cloned into the pHR lentiviral expression plasmid using inFusion cloning. Linear DNA fragments were produced by PCR using HiFi polymerase (Takara, 639298), followed by treatment with DpnI to remove template DNA. PCR products were then isolated through gel electrophoresis and purified using the Nucleospin gel purification kit (Takara Bio,740609.250). Linear DNA fragments were then ligated using inFusion assembly and amplified in Stellar competent *Escherichia* coli (Takara Bio, 636766). Plasmids were purified by miniprep (QIAGEN, 27104) and verified by whole-plasmid sequencing (Plasmidsaurus).

#### Cell line generation

Constructs were stably expressed in cells using lentiviral transduction. First, lentivirus was produced by co-transfecting HEK293T LX cells with pCMV-dR8.91, pMD2.G, and the expression plasmid of interest. 48 hr later, viral supernatants were collected and passed through a 0.45 µm filter. Cells were seeded at ∼40% confluency and transduced with lentivirus 24 hr later.

24hr post-seeding, culture medium was replaced with medium containing 10 µg/mL polybrene and 150–300 µL viral supernatant was added to cells. Cells were then cultured in virus-containing medium for 48 hr. Populations of cells co-expressing each construct were isolated using fluorescence-activated cell sorting on a Sony SH800S cell sorter. Sequentially bulk-sorted populations were collected for all experiments. We validated the cell lines used in this study (RPE, MCF10A) using STR profiling (Codes: sTRC4739,).

#### Tissue patterning

35mm glass bottom dish(CellvIs, D35-20-1.5-N) was coated with 10ug/ml human fibronectin (EMD Millipore, FC010) for 30 min at 37°C then washed three times with deionized water (DI). Surface of the dish was completely dried by nitrogen blowing. For the tissue seeding stencil, a 250μm thick PDMS membrane (Bisco HT-6240, Stockwell Elastomers) was cut by the Silhouette Cameo vinyl cutter. The stencil was treated with 2% pluronic F-127 (Invitrogen, P6866) solution diluted in PBS for 30 min at 37°C followed by three times wash with DI and drying with nitrogen. The dried stencil is attached to the fibronectin coated glass bottom dish.

Cells were washed once with PBS and treated TrypLE(Gibco, 12604-013) for 7 min at 37°C to be detached from the cell culture dish. The TrypLE treated cell solution was diluted with the culture medium and centrifuged for 3 min under 1500RPM. After the centrifugation, the supernatant was aspirated and the cell pellet was dissolved to the culture medium. The resuspended cell solution was carefully seeded into the stencil using micropipette. Concentration of the cell solution was aimed to be between 1.25E6 to 1.5E6 cells/ml. Seeding volume was determined by the empirical equation: seeding volume (μl) = stencil area(mm^2^) x conversion constant (0.44ul/mm^2^). To facilitate cell adherence, the cell seeded dish was incubated for 1 hr at 37°C before being flooded with the culture medium. 15 hr after flooding, the culture medium was exchanged to serum-free starvation medium consisting of DMEM/F12 (Gibco,11320033), 1% L-glutamine (Gibco,25030081), and 1% penicillin/streptomycin (Gibco,25030081). Imaging was performed 3 hr after the media exchange.

For OptoEGFR activity validation experiment (Fig. S1), cells were imaged on glass-bottom, black-walled 96-well plates (Cellvis, P96-1.5H-N) coated with fibronectin. Wells of 96-well plates were first incubated with 10 µg/mL fibronectin dissolved in PBS at 37°C for a minimum of 30 min. Cells were then seeded on glass-bottom 96-well plates at ∼40,000 cells/well 1 day prior to imaging. To increase adhesion, cell suspension is plated into 100 μL of media and then spun down in a tabletop centrifuge for 30 sec. After confirming the adhesion of cells an additional 100 μL of full media is added. The growth medium of cells was replaced with serum-free starvation medium 3 h prior to imaging.

#### Light guard generation for illumination pattern projection

The measurement of the scaling factor (length of physical pattern on light guard / length of projected illumination pattern on the dish) was measured with circular pattern light guard. Black plastic weighing boat (Heathrow Scientific, HS1423CC) was cut with a laser cutter to generate the light guard. The light guard was attached to the empty slot of the polarizer. Transmitted light source was turned on and the illumination pattern was focused by adjusting the height of the condenser turret. The image of the illumination pattern was captured and the diameter of the illuminated circle was measured with the ImageJ software. The scaling factor was calculated by dividing the diameter of the circular hole in the test light guard by the diameter of the circular illumination. The measured scaling factor for the Nikon Ti-2 system was 2.61. Based on the value, light guards with desired illumination patterns were designed and manufactured in the same way as our test light guard (**Fig. S2**).

#### Live-cell imaging

For small-scale patterning experiments (e.g., **Fig. 1**), imaging was performed on a Nikon Ti microscope with an iXon EM-CCD camera using a 20x objective. Patterned optogenetic stimuli were applied using a Mightex Polygon 4000 digital micromirror device (DMD) and an X-Cite XLED 450 nm light source. To prevent evaporation of media while imaging, 50 µL of mineral oil (VWR) was pipetted onto wells prior to mounting samples on the microscope. Optogenetic stimulation was achieved with the DMD set to a value of 75% and 200 um diameter ROI which resulted in a measured intensity at the objective lens of 65 mW/cm^2^.

For large-scale patterning experiments (e.g., **Fig. 2**), imaging was performed on a Nikon Eclipse Ti-2 microscope with a Qi-2 camera, RFP channel for RPE and MCF10A, using a 10x objective. Live-cell imaging was performed within the custom-made incubator box which maintains 37°C and supplies humidified 5% CO_2_ air flow. Images were captured every 10 minutes. To project illumination patterns, light guards were attached to the empty slot of the polarizer. The transmitted light source’s blue LED was used to apply 450 nm illumination to the tissue, with a measured intensity at the sample plane of 12 mW/cm^2^. The default illumination frequency for all figures was 5s / min except where otherwise indicated. Illumination frequency was adjusted to 4s / 2.5min for the high throughput assay in **Fig. 4C-G**. For live nuclear imaging, tissues were incubated in serum-free media with 10 μM Janelia Hoechst 646 for 1 h before imaging in the Cy5 channel.

#### Immunostaining

To quantify the 3D structure of the tissue (**Fig. 2E-G**), the tissues were fixed and stained with Nucblue formulation of the DAPI dye (Invitrogen, R37605) after the experiment. The tissues were treated with 4% PFA diluted in PBS for 45 minutes. 2 drops of Nucblue were added to each dish. Nuclei stained tissues were imaged by the W1 confocal unit, using the 20x objective.

#### Inhibition Assays

Cells were plated according to the protocols outlined above for the small and tissue scale conditions. After the 3-hour starvation period media containing the desired inhibitor concentration was added to each experimental well immediately before imaging. The following small molecule inhibitors were used:

| Name | Target | Concentration | Supplier/Cat # |
| --- | --- | --- | --- |
| Gefitinib | EGFR | 1 μM | MCE / HY-50895 |
| Cobimetinib | MEK1 | 1 μM | MCE / HY-13064 |
| MK2206 | AKT1/2/3 | 5 μM | Selleck Chem / S1078 |
| PI-103 | PI3K/ MTOR | 5 μM | MCE / HY-10115 |
| Gö6976 | pan-PKC | 5 μM | Selleck Chem / S7119 |
| TAPI-1 | ADAM17/ MMPs | 10 μM | MCE / HY-16657 |
| *para*-Nitro-Blebbistatin | non-muscle myosin II ATPases | 20 μM | Cayman Chem / 24171 |
| Y27632 | Rho kinase (ROCK) | 5 μM | Tocris / 1254 |

#### Immunoblotting analysis

To collect cell lysate, cells were cultured in 10mm tissue culture dishes, washed with PBS and lysed in RIPA buffer (ThermoFisher, 89900). Cell scrapers were used to separate adherent cells from the culture dish, and immediately placed on ice inside 1.5mL Eppendorf tubes. Lysates were centrifuged at 13,000 RPM at 4°C for 10 min, the supernatant was collected and the pellet was discarded. NuPAGE LDS sample buffer (Invitrogen, NP0007) was added to each sample before being heated at 95°C for 10 min and then placed on ice. Sample proteins were then separated via SDS-PAGE and transferred to nitrocellulose membranes using the iBlot 2 Gel Transfer (Invitrogen, IB21001). Membranes were blocked with Odyssey Blocking Buffer (LICOR 927-60001) for 1 h at room temperature preceding primary antibody incubation using a 1:1 mixture of Odyssey Blocking Buffer and TBST (ThermoFisher, J77500.K8) diluted to 1x concentration at 4°C overnight. The following primary antibodies were used: Phospho-p44/42 MAPK (Erk1/2) XP® (CST,4370), p44/42 Erk1/2 (CST 4696), β-Actin (CST,3700), and GAPDH (CST,D4C6R). The following secondary antibodies were used: IRDye® 800CW Goat anti-Rabbit IgG (Licor,926-32211) and IRDye® 680RD Goat anti-Mouse IgG (Licor,926-68070). Immunofluorescence imaging was conducted using the Licor Odyssey Clx system.

### Quantification and statistical analysis

#### Tissue migration analysis

Local velocity vector field of the tissue was generated by particle image velocimetry (PIV). PIVLab MATLAB plugin was used for this analysis^71^. Pass 1 and 2 window sizes were assigned as 200 pixels and 100 pixels for the timelapse image sequences captured with the 10x objective. Overlap between box was 50%. Further analysis was conducted after deducing radial velocity from the velocity vector field. Radial velocity was calculated by multiplying the speed to the cosine of angle difference between velocity vector and vector pointing toward the center of the illumination (**Fig. 2**; **Fig. S3**) or the tissue (**Fig. 3**). Peak radial velocity was defined as the peak value of radial velocity of the entire tissue. Migration zone width was measured by the distance from the illumination boundary to the point where the radial velocity value drops to the threshold value (1 μm/h), a sufficiently high value that is never attained by unilluminated tissue.

#### Pseudo density analysis

Confocal stack of nuclei stained tissue was processed by the sum Z-projection function in ImageJ. Radial average intensity from the illumination center was calculated with the ImageJ plugin Radial Profile Angle. The radial intensity data was binned with a 50μm binning window. Pseudo density was deduced by normalizing the fluorescence intensity by the average intensity of the control tissue.

#### Tissue height analysis

Confocal stack of nuclei stained tissue was radially reslice via reslice function in ImageJ. The radial reslice stack was further processed with the sum Z-projection function in ImageJ. The obtained merged XZ slice image was segmented with the threshold function in ImageJ. Local tissue height was measured with this binary image and was binned with a 50μm binning window.

#### Radius and edge expansion speed analysis

RPE tissue was segmented using Bernsen method in auto local threshold function in ImageJ. Area of the segmented tissue image was measured by the regionprop function in MATLAB. Radius of the tissue was calculated by fitting the tissue area to the circle. Edge expansion speed was deduced as the speed of radius increment.

#### Cell density analysis

Cy5 nuclei channel images were segmented by StarDist ImageJ plugin^72,73^. Centroid of each nucleus was calculated with the regionprop function in MATLAB. Local cell density was measured by (Number of centroids in the ROI)/(Area of ROI). Dimension of ROI was 100 pixel x 100 pixel square and 50% overlap between the adjacent ROIs.

#### Single cell analysis

The Cy5 channel nucleus image stack was tracked via the TrackMate ImageJ plugin. Persistence and speed were calculated using a custom MATLAB script.

#### Statistical test

Mann Whitney U-test was applied for statistical tests to compare differences between two groups. For the group size below 50, we used t-test analysis in Prism. For the group size above 50, we random-sampled 50 observations with replacement for each data group. P-value was calculated from these subsets. Mean P-value of 50 repeats of this process was used to decide statistical significance of the difference between two groups. Custom MATLAB script was used for this process.

#### Mathematical modeling

We constructed a simple mathematical model to obtain qualitative insights for how tissues might flow under patterned light inputs. Our model consists of a continuous variable *c(x,y)* representing the density of tissue at each position in 2D space. It also incorporates an arbitrarily-drawn light input *u(x,y)* that takes binary values (1 for illumination at that position; 0 for darkness). The model incorporates two cellular processes: a diffusion term (with diffusion constant *D*) to represent tissue spreading over time and a flux term at boundaries of the binary illumination input in the direction of the light with rate *k*. We simulated this partial differential equation system by discretizing the *x* and *y* coordinates into 101 bins and simulating the resulting 10,201 element ordinary differential equation system, where each element was defined as:

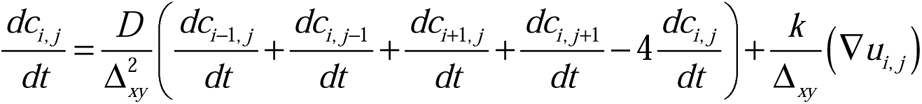

where ∇*u_i_* _, *j*_ is related to the gradient of the light stimulus, and is defined as 1 for elements where the input *u* changes from 0 to 1 and -1 for elements where the input changes from 1 to 0 for pairs of elements along the *x* or *y* direction and Δ*_xy_* is the length scale associated with each discretized spatial element (e.g., the total length scale of simulation divided by 101). MATLAB code implementing the model is available at the Github (https://github.com/toettchlab/Suh-Thornton2024).

To obtain approximate values for the parameters *D* and *k*, we simulated the 1 mm diameter tissue densification pattern of **Fig. 2G** (see **Fig. S4**) to qualitatively match the length and time scale of tissue movement into the illuminated region, which led us to values *D*=50 μm^2^/min and *k*=0.3 min^-1^. We then simulated the complex pattern of **Fig. 6** using the same parameters.

## Acknowledgments

The authors thank all members of the Toettcher and Cohen labs, particularly Sabrina Solley and Beatrice Ramm for help throughout the project. Figure illustrations were created in part using Biorender. This work was supported by NIH grant T32GM007388 and a Janssen Scholars of Oncology Diversity Engagement Program (SODEP) Award (to R.H.T.); NIH grant R01GM144362 (to J.E.T.), and funding from the Omenn-Darling Bioengineering Institute (to J.E.T. and D.J.C.).

## Author contributions

Conceptualization, K.S., R.H.T., D.J.C., P.E.F., J.E.T.; Methodology, K.S., R.H.T., P.E.F., D.J.C., J.E.T.; Investigation, K.S., R.H.T., P.E.F.; Funding, R.H.T., D.J.C., J.E.T.; Writing and Editing, K.S., R.H.T., D.J.C., J.E.T.; Supervision, D.J.C., J.E.T.

