## Supplementary Information for "Large-scale control over collective cell migration using light-controlled epidermal growth factor receptors"

### Supplementary Figures

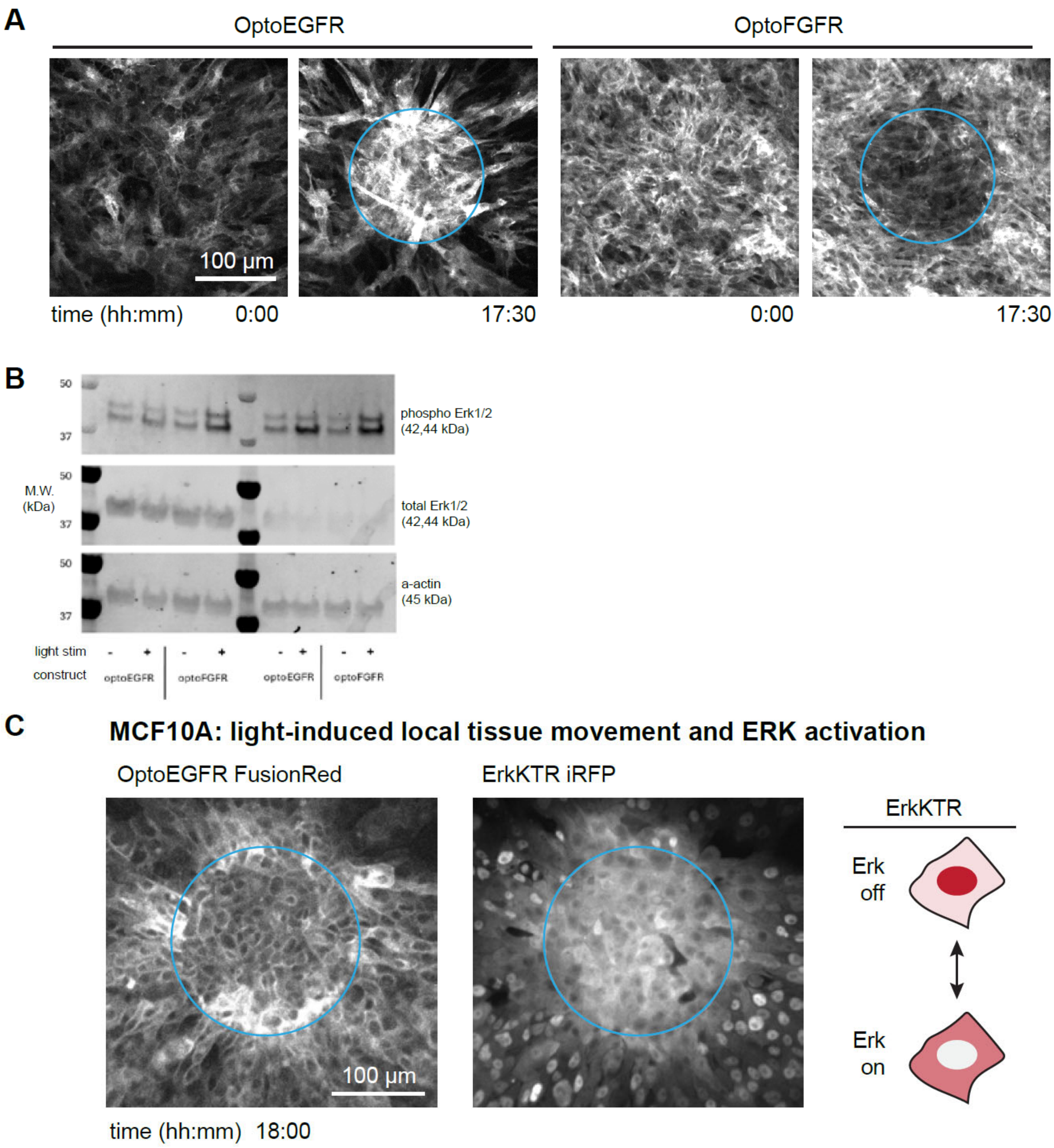

**Figure S1. Light-induced cell motility is a specific feature of OptoEGFR activation and generalizes to multiple cellular contexts.** (**A**) Representative fields of view showing OptoEGFR-induced tissue movement toward regions of illumination and OptoFGFR exclusion from illumination regions. (**B**) Western blot of independently treated OptoEGFR and OptoFGFR cell lysates in light and dark, showing phospho-ERK, total ERK and actin as a loading control. Both OptoEGFR and OptoFGFR show similar levels of ERK phosphorylation upon illumination, indicating similar degrees of light-induced activation. (**C**) OptoEGFR stimulation of cell migration in MCF10A cells also expressing the ErkKTR-iRFP, which indicates ERK activity by a shift from nuclear to cytosolic localization. Light induces local ERK activation within the illuminated region as well as tissue movement toward the light input on a similar timescale to what is observed in RPE-1 OptoEGFR cells. Related to Figure 1.

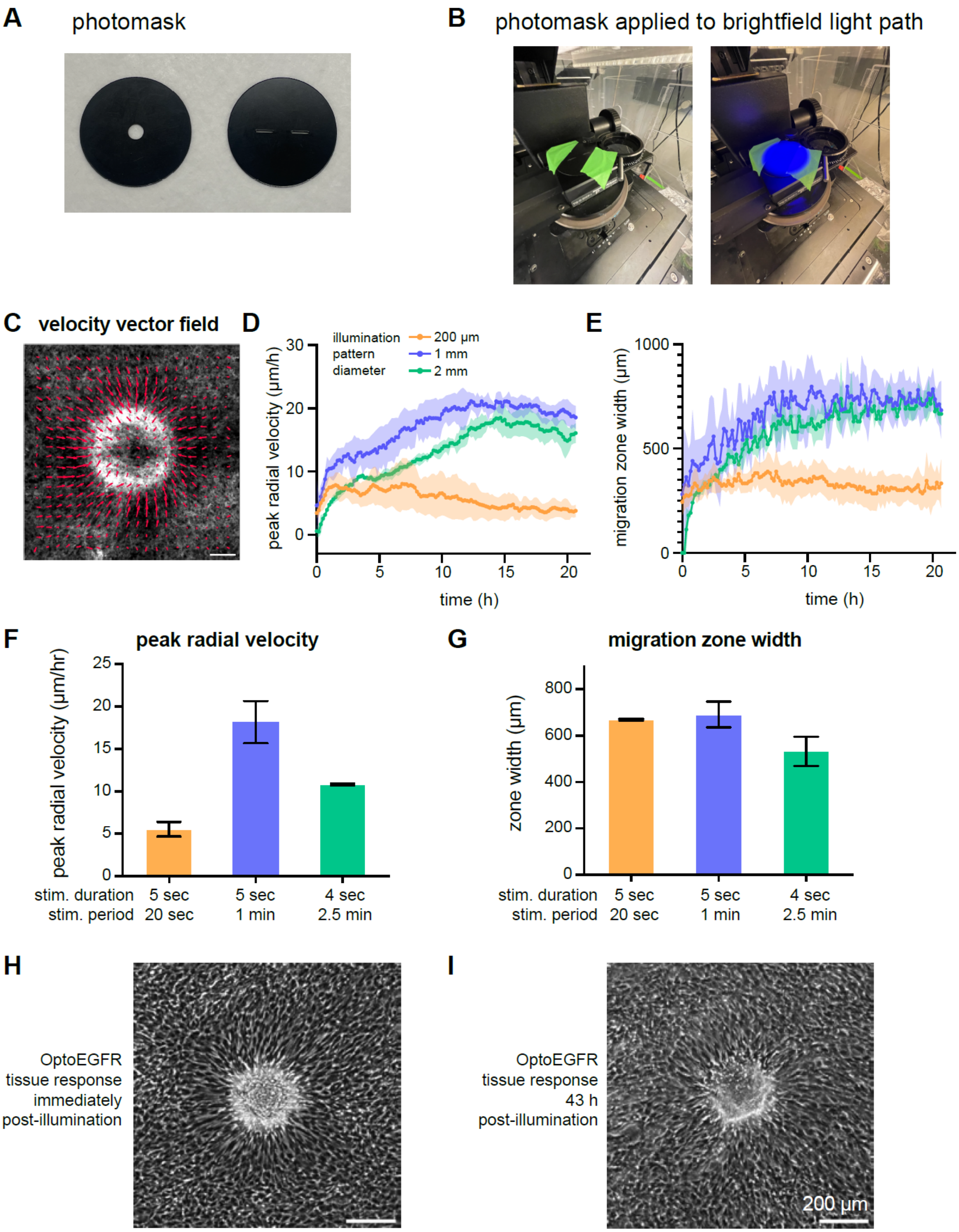
A
photomask
B
photomask applied to brightfield light path
C
velocity vector field
D
illumination pattern diameter
200 µm
1 mm
2 mm
peak radial velocity (µm/h)
time (h)
E
migration zone width (µm)
time (h)
F
peak radial velocity
peak radial velocity (µm/hr)
stim. duration
stim. period
5 sec
20 sec
5 sec
1 min
4 sec
2.5 min
G
migration zone width
zone width (µm)
stim. duration
stim. period
5 sec
20 sec
5 sec
1 min
4 sec
2.5 min
H
OptoEGFR tissue response immediately post-illumination
I
OptoEGFR tissue response 43 h post-illumination
200 µm

**Figure S2. Large-scale optical illumination drives tissue flows at a millimeter scale.** (A-B) Experimental setup for illumination light shield and DMD. The laser-cut photomask (shown in A) is placed on top of the LED-based transmitted light turret, which is used to deliver 450 nm blue light. (C) Velocity vector field for tissue flows near the 1 mm diameter illumination pattern. (Scale bar: 300 µm). (D-E) Maximum radial velocity (in D) and migration zone width (in E) plotted over time for 200 µm, 1 mm, and 2 mm illumination patterns. For larger patterns, tissue flow is sustained over the timecourse with a peak at ~15 h; for the 200 µm pattern, tissue movement is primarily observed during the first ~10 h with a peak at ~3 h. (F-G) Peak radial velocity and migration zone width shown as a function of illumination dose. Light inputs of identical intensity were delivered for 5 sec every 20 sec, for 5 sec every 1 min, or for 4 sec every 2.5 min. The OptoDroplet light-responsive module in OptoEGFR has a dark-state reversion time of ~2-3 min, so each stimulus type is expected to drive different intermediate activity levels that are sustained over time. (H-I) Persistent tissue deformation observed even 43 h after removal of the optogenetic stimulus. Brightfield images are shown immediately after applying a 36 h stimulus or after waiting an additional 43 h after stimulation. Related to Figure 2.

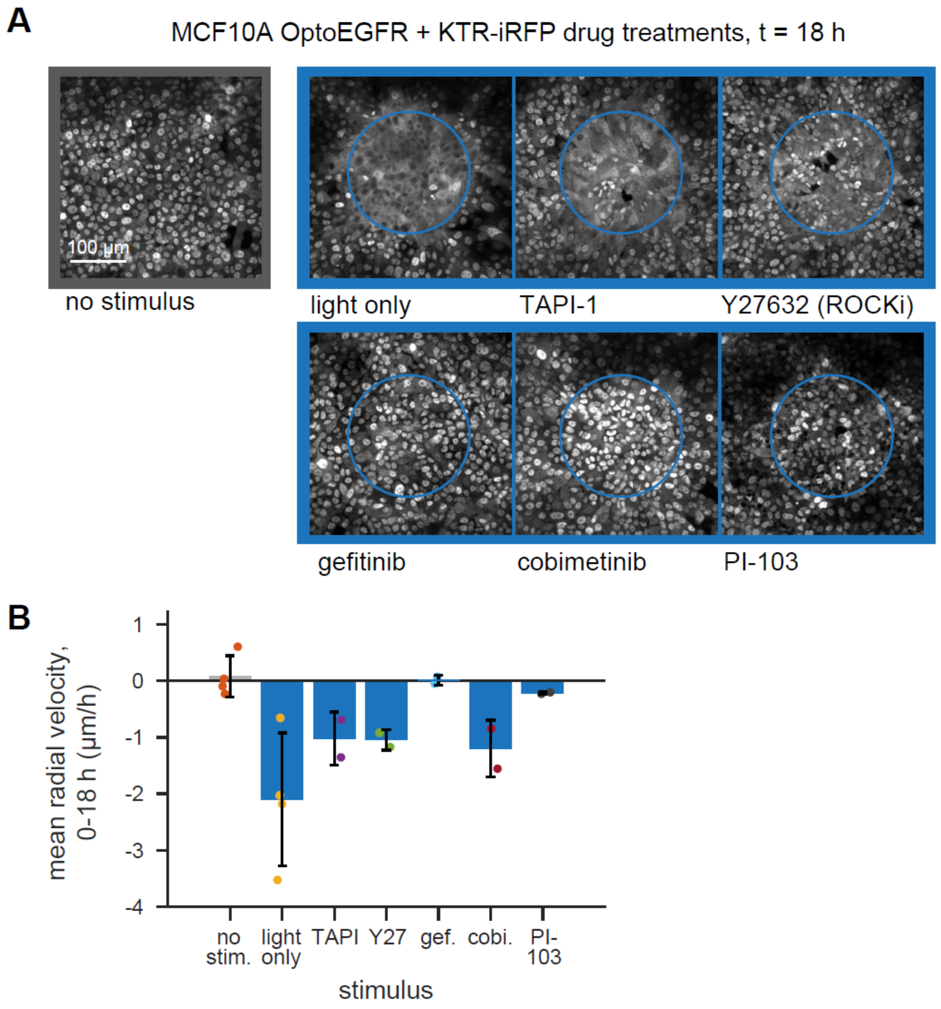


**Figure S3. Effect of chemical inhibitors on light-induced migration in MCF10A cells.** (A) Representative images of MCF10A OptoEGFR and ErkKTR-iRFP expressing cell 18 h after treatment with no input, a local pattern of 450 nm light ("light only"), the matrix metalloproteinase inhibitor TAPI-1, the Rho kinase inhibitor Y27632 which is expected to inhibit myosin activation, the EGFR inhibitor gefinitib, the MEK inhibitor cobimetinib, or the PI 3-kinase inhibitor PI-103. ErkKTR-iRFP fluorescence imaging shown. (B) Quantification of mean radial tissue velocity over the 18 h timecourse in each stimulus condition shown in A. Related to Figure 4.

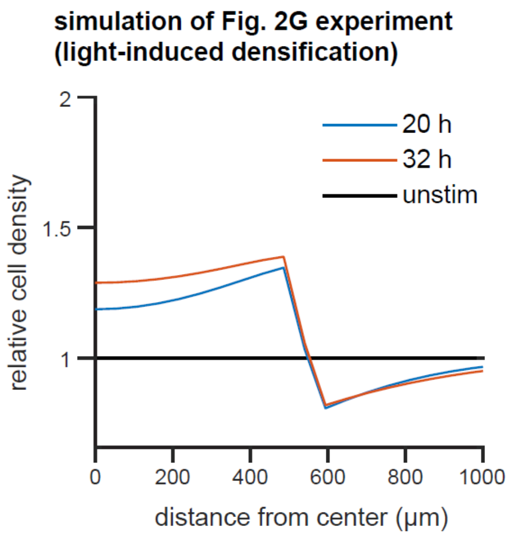


**Figure S4. Model simulation of tissue convergence from Fig. 2G.** A 1 mm diameter (500 µm radius) circular pattern of illumination was simulated in the model and the density of tissue was determined after 20 and 32 h simulation time. The depletion zone and peak of cell density at the illumination boundary are evident from the simulation, which only incorporates boundary-driven tissue flux into the illuminated region and diffusion-based tissue spreading. Related to Figure 5.

#### Supplementary Videos

**Video S1.** Light-induced movement of RPE-1 OptoEGFR cells towards a region of local illumination (blue circle). Scale bar: 100 µm. Time shows hh:mm after stimulation. Related to Figure 1.

**Video S2.** Light-induced movement of RPE-1 OptoEGFR cells outward from a tissue boundary after removal of a barrier and local illumination (blue rectangle). Scale bar: 100 µm. Time shows hh:mm after stimulation. Related to Figure 1.

**Video S3.** Light-induced movement of MCF10A OptoEGFR ErkKTR-iRFP cells towards a region of local illumination (blue circle). Scale bar: 100 µm. Time shows hh:mm after stimulation. Related to Figure 1.

**Video S4.** Millimeter-scale cell responses of RPE-1 OptoEGFR cells to illumination patterns of varying sizes. Scale bar: 1 mm. Time shows hh:mm after stimulation. Related to Figure 2.

**Video S5.** Tissue outgrowth of RPE-1 OptoEGFR cells after release from confinement to a circular pattern in the presence or absence of global 450 nm illumination. Scale bar: 500 µm. Time shows hh:mm after stimulation. Related to Figure 3.

**Video S6.** Tissue outgrowth of RPE-1 OptoEGFR tissues initially separated by a 300 µm gap, with the right-hand tissue exposed to 450 nm light throughout the experiment. Scale bar: 500 µm. Time shows hh:mm after stimulation. Related to Figure 4.

**Video S7.** Responses of RPE-1 OptoEGFR cells to a 200 µm diameter local illumination pattern in the presence of various chemical inhibitors. Scale bar: 500 µm. Time shows hh:mm after stimulation. Related to Figure 4.

**Video S8.** Responses of MCF10A ErkKTR-iRFP cells that also express OptoEGFR cells (left) or OptoSOS (right) to local light stimulation. ErkKTR-iRFP fluorescent images are shown, which demonstrate that both constructs induce similar ERK activity in the illuminated region but only OptoEGFR elicits a migratory response. Scale bar: 100 µm. Time shows hh:mm after stimulation. Related to Figure 4.

**Video S9.** Complex tissue patterning using optogenetic stimulation of RPE-1 OptoEGFR cells. Scale bar: 1 mm. Related to Figure 5.
